# Lipid metabolic stress in development defines which genetically-susceptible DYT-*TOR1A* mice develop disease

**DOI:** 10.1101/2020.03.18.997247

**Authors:** Ana Cascalho, Joyce Foroozandeh, Stef Rous, Natalia Martínez Vizcaíno, Sandra F. Gallego, Rose E. Goodchild

## Abstract

There has been enormous progress defining the genetic landscape of disease. However, genotypes rarely fully predict neurological phenotypes, and we rarely understand why. *TOR1A* +/Δgag that causes dystonia with ~30% penetrance is a classic case. Here we show, in inbred mice, that +/Δgag affects embryonic brain lipid metabolism with sex-skewed reduced penetrance. Penetrance is affected by environmental context, including maternal diet. The lipid metabolic defect resolves during post-natal development. Nevertheless, we discover dystonia-like symptoms in ~30% of juvenile female *Tor1a*^+/Δgag^ mice, and prevent these symptoms by genetically suppressing abnormal lipid metabolism. We conclude that *Tor1a*^+/Δgag^ embryos poorly buffer metabolic stress *in utero*, resulting in a period of abnormal metabolism that hardwires the brain for dystonia in later life. The data show unexpected and profound impacts of sex, and thus highlight the importance of examining male and female animal models of disease.

**Significance Statement:** The genetic landscape of neurological disease is relatively well mapped. However, we typically cannot explain why some mutations only cause disease in a subset of individuals. A classic case is DYT-*TOR1A* dystonia that only develops in 30% of *TOR1A*^+/Δgag^ carriers. We now find that ~30% of inbred female *Tor1a*^+/Δgag^ mice develop abnormal brain lipid metabolism as embryos, while males are spared. The percentage is affected by maternal diet. Further, this period of abnormal lipid metabolism causes dystonia-like symptoms in juvenile mice. These data show how an environmentally-sensitive event of development defines which genetically-susceptible individuals develop disease in later life. They also highlight the importance of examining male and female animal models of disease.

## Introduction

Neurogenetics has associated hundreds of mutations with neurological disease. However, genotype does not always predict phenotype (Cooper et al., 2013). This includes the phenomenon of reduced penetrance where a “causative” mutation produces symptoms in some individuals, while others escape disease. It is thought this arises when genetic and/or environmental modifiers suppress or enhance pathological process(es) triggered by the mutation. There is also the hypothesis that susceptibility or resistance to *in utero* stresses is key for some disorders. This “developmental origins of health and disease” hypothesis has strong support from human epidemiology (Gluckman et al., 2010). It also applies to complex conditions like autism or schizophrenia that have a major genetic component but, for poorly defined reasons, even monozygotic siblings variably develop symptoms (Kim and Leventhal, 2015; Nestler et al., 2016; Singh et al., 2004). A major challenge is the lack of experimental systems that can explicitly test these concepts.

The dominant childhood-onset movement disorder DYT-*TOR1A* dystonia (OMIM #128100) is a classic reduced penetrance disease. It is characterized by involuntary twisting movements and abnormal postures that appear at a median age of 8-11 years in 30-40% of individuals carrying a 3-base pair deletion in *TOR1A* (Δgag) (Ozelius and Lubarr, 1993). There is no clear answer for why only some *TOR1A*^+/Δgag^ individuals develop dystonia (Martino et al., 2013). A protective polymorphism exists in *TOR1A*, but is too rare to explain penetrance at the population level (Frederic et al., 2009; Kamm et al., 2008). The frequency of DYT-*TOR1A* dystonia is also similar between ethnicities and geographical regions (Kramer et al., 1994; Yang et al., 2009). The disease is incurable, and its underlying neurobiology remains mysterious, including that there is no overt neuropathology that explains symptoms.

Information on the molecular function of *TOR1A* / TorsinA provides an alternative route to understand DYT-*TOR1A* dystonia. Torsin proteins have emerged as critical regulators of cellular lipid metabolism. They especially affect phosphatidic acid (PA) metabolism to diacylglycerol (DAG) mediated by the Lipin family of enzymes (Figure 1A). PA and DAG are signaling lipids that activate a number of pathways (Wang et al., 2006; Young et al., 2010). They impact membrane dynamics (Zhukovsky et al., 2019) and are precursors for competing lipid metabolic pathways (Craddock et al., 2015; Yang et al., 2020). The link to Torsin includes that fly *dTorsin* regulates PA and DAG levels, and *dTorsin^KO^* larvae have excess triglyceride and lipid droplet accumulation (Grillet et al., 2016). *Tor1a* deletion from mouse liver also causes a fatty liver phenotype of excess triglycerides and lipid droplets (Shin et al., 2019), and the brains of *Tor1a* knock-out mice have ~3-fold higher Lipin-mediated PA to DAG conversion (Cascalho et al., 2020). Moreover, the molecular function is conserved in humans: iPSC-derived neurons from *TOR1A*^+/Δgag^ dystonia patients have ~1.5-fold higher Lipin-mediated PA to DAG conversion (Cascalho et al., 2020).

**Figure 1.**
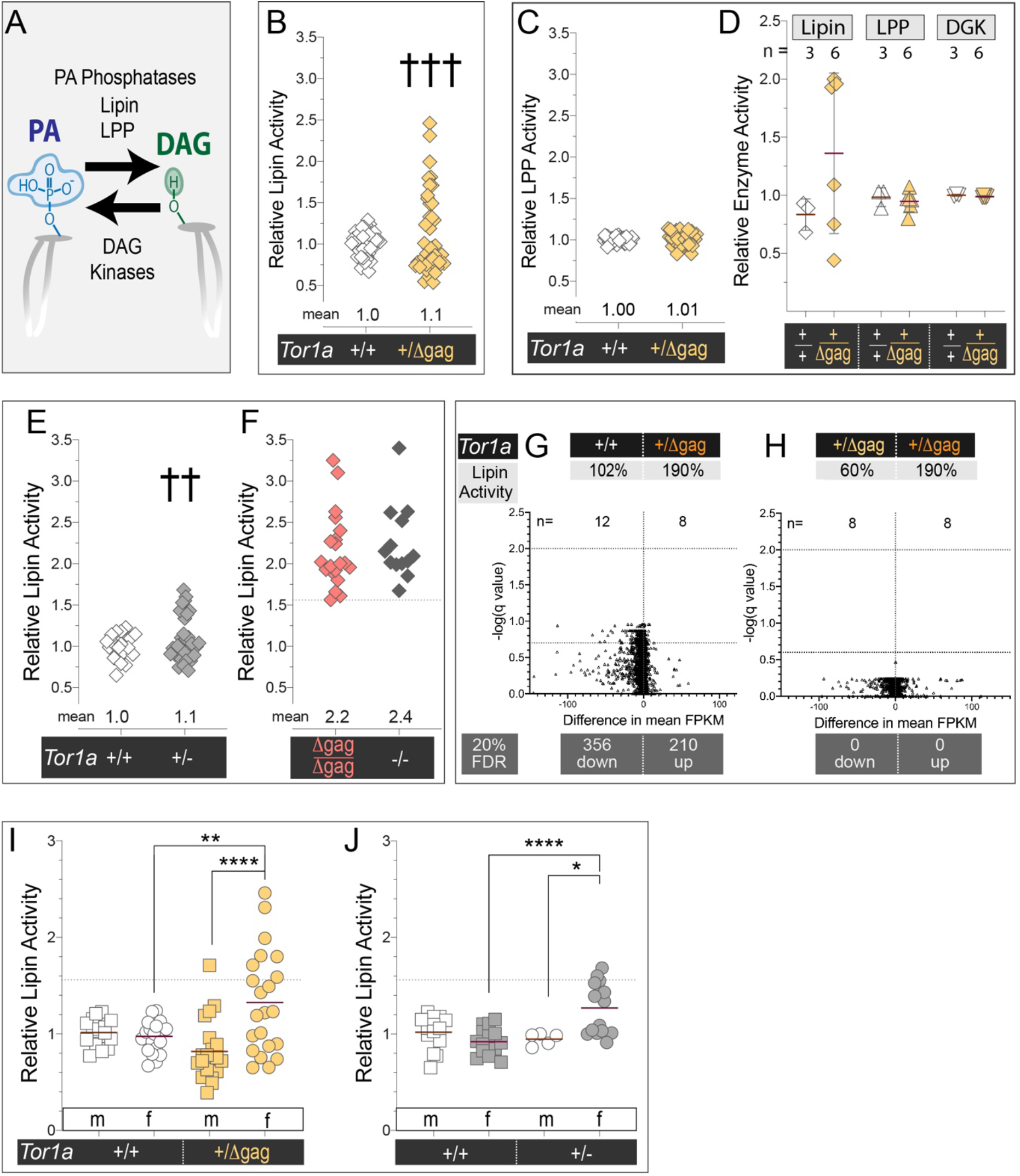
*Tor1a*^+/Δgag^ embryos display excess brain Lipin activity with sex-skewed reduced penetrance. A) Lipin, LPP and DAG kinases interconvert PA and DAG. B) Lipin activity in brain homogenates from individual E18.5 embryos. ††† indicates significantly non-normal distribution (p < 0.001, Kolmogorov-Smirnov test). Data is presented relative to the *Tor1a*^+/+^ mean. C) LPP activity in E18.5 brain homogenates. D) Lipin, LPP and DAG Kinase (DGK) activities in E18.5 brain homogenates. Bars show mean +/− SD. E & F) as (B); †† indicates significantly non-normal distribution (p < 0.01, Kolmogorov-Smirnov test). Dotted line shows the lowest level of Lipin activity in the brain of homozygous *Tor1a* mutant embryos. G & H) Volcano plots show multiple T-Test comparisons of FPKM (Fragments Per Kilobase of transcript per Million mapped reads) values of (G) *Tor1a*^+/+^ versus *Tor1a*^+/Δgag^ (17777 genes) and (H) *Tor1a*^+/Δgag^ embryos with low or elevated Lipin activity (17776 genes). Dotted lines highlight 20% FDR. Percentage values are the mean Lipin activity of each group compared to *Tor1a*^+/+^. “Down” and “up” refer to the number of differentially expressed genes according to a 20% FDR. I & J) Lipin activity in E18.5 brain homogenates: “m” male, “f” female. Dotted line defines the lowest value of Lipin activity in brain homogenates from homozygous *Tor1a* mutant embryos. (I) Two-Way ANOVA detects a significant effect of sex (p = 0.003), and interaction between sex and *Tor1a* genotype (p = 0.0006). (J) Two-Way ANOVA detects a significant effect of sex (p = 0.02), and interaction between sex and *Tor1a* genotype (p = 0.001).

It has been challenging to address if lipid metabolism contributes to DYT-*TOR1A* dystonia because +/Δgag does not appear to induce disease in experimental models (Tanabe et al., 2012; Ulug et al., 2011). Nevertheless, it is intriguing to consider the many environmental factors that regulate lipid metabolism and might mitigate or amplify the impact of *TOR1A*^+/Δgag^. This includes that the Lipin1 enzyme is directly phosphorylated and inhibited by mTORC1 (Huffman et al., 2002; MacVicar et al., 2019; Peterson et al., 2011; Romani et al., 2019). We therefore examine the relationship between +/Δgag and lipid metabolism, as well as addressing the reduced penetrance of DYT-*TOR1A* dystonia.

## Results

### +/Δgag affects brain Lipin activity with sex-skewed reduced penetrance

We measured Lipin activity via the production of fluorescent nitrobenzoxadiazole-DAG (NBD-DAG) from NBD-PA added to brain homogenates (Supplemental Figure 1A). We examined embryonic day 18.5 (E18.5) *Tor1a*^+/+^ and *Tor1a*^+/Δgag^ brains, which is the developmental stage when *Tor1a* knock-out mice have highly elevated Lipin activity (Cascalho et al., 2020). Lipin-mediated DAG production was similar between individual *Tor1a*^+/+^ embryos, and these values were normally distributed (Figure 1B). Surprisingly, mean Lipin activity only mildly differed between *Tor1a*^+/+^ and *Tor1a*^+/Δgag^ (p = 0.03, Kolmogorov-Smirnov non-parametric test). However, more strikingly, values from *Tor1a*^+/Δgag^ embryos were not normally distributed, and ~16% had higher Lipin activity than any *Tor1a*^+/+^ embryo (Figure 1B).

The *Tor1a*^+/Δgag^ mouse has been backcrossed with C57BL6/J for > 20 generations, and genetic SNP analysis of six *Tor1a*^+/Δgag^ animals defined these as 99.9% C57BL/6. Thus, there is no obvious genetic explanation for variability between individual animals. We validated that varied DAG accumulation indeed represented variable Lipin activity. The assay was linear between 0.25 and 3-fold of the activity in *Tor1a*^+/+^ homogenates (Supplemental Figure 1B), and it detected a dose-dependent reduction between *Lipin1*^+/+^ *Lipin1*^+/−^ and *Lipin1*^−/−^ brains (Supplemental Figure 1C). Lipid Phosphate Phosphatase (LPP) enzymes perform PA to DAG conversion. LPP activity was similar between homogenates of *Tor1a*^+/+^ and *Tor1a*^+/Δgag^ brains (Figure 1C). We directly compared PA to DAG metabolic reactions of several *Tor1a*^+/+^ and *Tor1a*^+/Δgag^ brains. This re-confirmed the *Tor1a*^+/Δgag^ group had variable magnesium-dependent PA to DAG conversion (Lipin; Figure 1D left columns), normal magnesium-independent conversion (LPP; Figure 1D middle columns), and normal DAG kinase conversion of NBD-DAG to NBD-PA (Figure 1D, right columns). We therefore conclude that variable DAG accumulation derives from variable Lipin activity.

Genetics and biochemistry show that Δgag inhibits *Tor1a* (Demircioglu et al., 2016; Goodchild et al., 2005; Zhao et al., 2013). We turned to *Tor1a* knock-out mice as an independently generated and inbred line to further examine the effect of *Tor1a* genotype on lipid metabolism. Individual *Tor1a*^+/+^ embryos from *Tor1a*^+/−^ crosses had normally distributed Lipin activity (Figure 1E). In contrast, the *Tor1a*^+/−^ population was a) not normally distributed, and b) contained a subpopulation that had higher brain Lipin activity than all *Tor1a*^+/+^ littermates (Figure 1E). *Tor1a*^Δgag/Δgag^ and *Tor1a*^−/−^ die as neonates concomitant with mean brain Lipin activity > 3-fold above normal (Figure 1F; (Cascalho et al., 2020)). We used the lowest values from homozygous *Tor1a* mice to define “pathogenic” Lipin activity. This was ~1.5 fold above the wild-type mean. 3% of *Tor1a*^+/−^ and 16% of *Tor1a*^+/Δgag^ embryos had Lipin activity at or above this level.

We sought factors that explained the variation of heterozygous *Tor1a* embryos. TorsinA and homologous TorsinB proteins were similar between brains with different Lipin activity (Supplemental Figure 2A). There was no relationship between Lipin activity and *Lipin1* and *Lipin2* gene expression (Supplemental Figure 2B – C), nor for genes that regulate Lipins or Torsins (Supplemental Figure 2D - G). We compared genome-wide RNAseq profiles between *Tor1a*^+/+^ and *Tor1a*^+/Δgag^ brains, as well as between a *Tor1a*^+/Δgag^ group with low Lipin activity and a *Tor1a*^+/Δgag^ group with high Lipin activity (Supplemental Figure 2H). A low-stringency 20% false discovery rate (FDR) defined ~600 differentially expressed genes between *Tor1a*^+/+^ and *Tor1a*^+/Δgag^ (Figure 1G; Supplemental TABLE 1). However, no genes were differentially expressed between *Tor1a*^+/Δgag^ groups with different Lipin phenotypes (Figure 1H).

The consistent gene expression profiles again argue against a genetic origin for the phenotypic variability. We therefore considered factors that drive variability. *Tor1a*^+/Δgag^ embryos with normal and high Lipin activity coexisted in litters, and there was no clear relationship with litter size (Supplemental Figure 3A). Lipin activity was similarly variable for *Tor1a*^+/Δgag^ embryos derived from *Tor1a*^+/Δgag^ or wild-type C57BL/6J dams. We considered the role of sex. There was no difference in brain Lipin activity between male and female *Tor1a*^+/+^ embryos. However, sex strongly influenced Lipin activity in the *Tor1a*^+/Δgag^ population. Female *Tor1a*^+/Δgag^ brains had higher Lipin activity than male (Figure 1I). Further, all but one of the *Tor1a*^+/Δgag^ embryos with “pathogenic” Lipin hyperactivity were female. This amounted to 35% of female *Tor1a*^+/Δgag^ embryos (Figure 1I). We also examined the effect of sex on *Tor1a*^+/−^ embryos. Again, female *Tor1a*^+/−^ had higher brain Lipin activity than male. Further, the entire *Tor1a*^+/−^ subpopulation with higher than normal values were female (Figure 1J).

### Environment defines whether +/Δgag deregulates lipid metabolism

Sex influences development via X- and Y-chromosome gene expression in the embryo, as well as in the extra-embryonic placenta that, in turn, controls embryo exposure to nutrients and stresses (Bale, 2016; McNairn et al., 2019; Werner et al., 2017). Male testosterone signaling also commences *in utero*, where it variably affects female littermates depending on their intrauterine proximity to males (vom Saal and Bronson, 1980). We measured Lipin activity in E13.5 embryonic brains when there is minimal testosterone production (O’Shaughnessy et al., 1998). This again identified a proportion of female *Tor1a*^+/Δgag^ with excess brain Lipin activity while males were unaffected (Figure 2A), pointing away from a sex hormone explanation.

**Figure 2.**
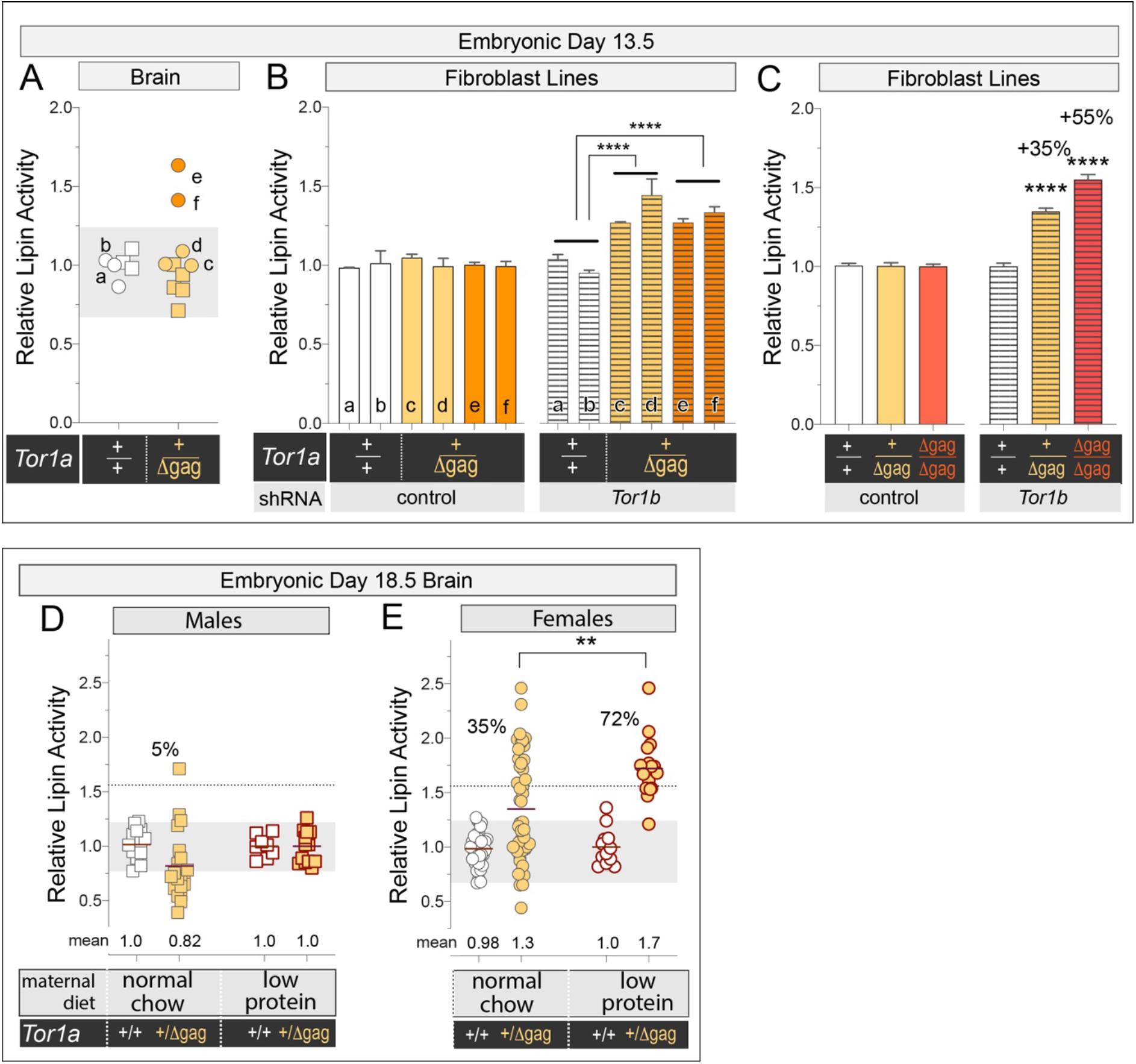
Environment modifies whether *Tor1a*^+/Δgag^ affects lipid metabolism. A) Lipin activity in E13.5 brain homogenates. Circles show females, squares show males. Shaded zone shows the highest and lowest values from the *Tor1a*^+/+^ population. B) Lipin activity of MEFs derived from the embryos indicated in (A), and electroporated with empty or *Tor1b* shRNA vectors. Bars show the mean +/− SEM from duplicate electroporations presented relative to the mean of *Tor1a*^+/+^ with empty vector. Two-way ANOVA detects a significant effect of *Tor1a* genotype (p < 0.0001), *Tor1b* knockdown (p < 0.0001), and interaction between *Tor1a* and *Tor1b* (p = 0.0002). There is no difference between the four *Tor1a*^+/Δgag^ cultures (p = 0.55). C) Lipin activity in MEF homogenates electroporated with empty or *Tor1b* shRNA vectors. Two-way ANOVA detects significant effect of *Tor1a* genotype, *Tor1b* knockdown, and interaction between *Tor1a* genotype and *Tor1b* knockdown (p < 0.0001). **** indicates significant difference compared to *Tor1a*^+/+^. Values show the % increase in Lipin activity compared to *Tor1a*^+/+^. D & E) Lipin activity in E18.5 brain homogenates. Dotted line and percentages refer to the number of animals with Lipin activity in the range defined by homozygous *Tor1a* mutants. (E) Two-Way ANOVA detects a significant effect of *Tor1a* genotype (p < 0.0001), maternal nutrition (p = 0.01), and interaction between *Tor1a* genotype and maternal nutrition (p = 0.02).

We asked whether *Tor1a*^+/Δgag^ acts with reduced penetrance under standardized *in vitro* conditions. We prepared mouse embryonic fibroblasts (MEF) lines from female *Tor1a*^+/+^ and *Tor1a*^+/Δgag^ that had varied brain Lipin activity (Figure 2A). MEFs co-express TorsinB with TorsinA, which protects against *Tor1a* mutations (Cascalho et al., 2020; Kim et al., 2010). Consistently, baseline Lipin activity was similar across all MEF lines (Figure 2B, left bars). We induced a similar degree of *Tor1b* knockdown across the lines (Supplemental data Figure 3B). This consistently increased Lipin activity in the *Tor1a*^+/Δgag^ lines, showing that *in vivo* variability of *Tor1a*^+/Δgag^ is not maintained *in vitro* (Figure 2B, right bars). We examined an allelic series of *Tor1a*^+/+^, *Tor1a*^+/Δgag^, and *Tor1a*^Δgag/Δgag^ MEFs. This defined the ~1.3-fold Lipin hyperactivity of *Tor1a*^+/Δgag^ lines at approximately half way between *Tor1a*^+/+^ and *Tor1a*^Δgag/Δgag^ (Figure 2C; Supplemental data Figure 3C). We therefore conclude that +/Δgag is a haploinsufficient allele in a standard *in vitro* cell culture context.

**Figure 3.**
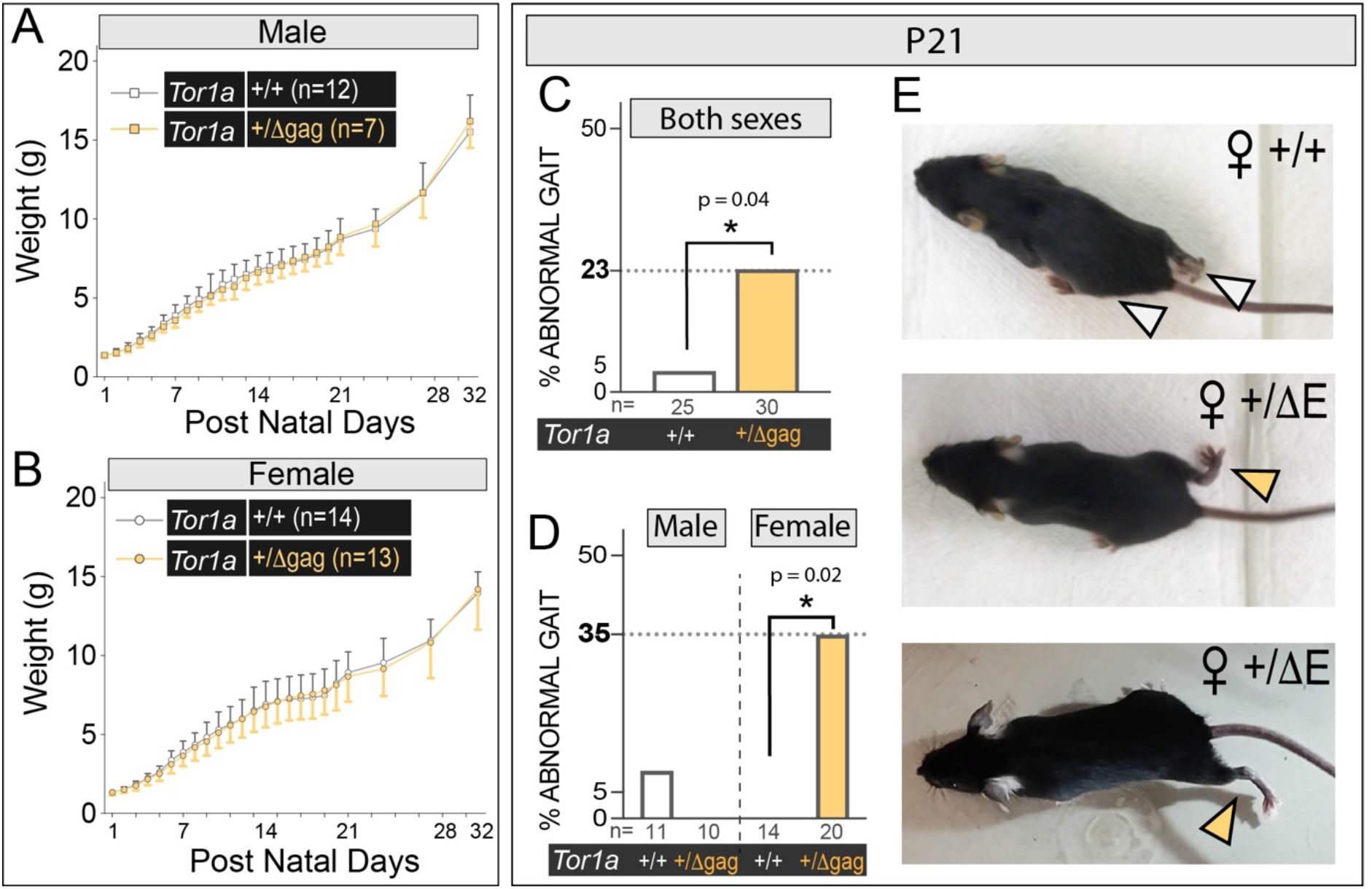
A dystonia-like movement disorder in ~30% of female *Tor1a*^+/Δgag^ mice. A & B) *Tor1a*^+/Δgag^ has no effect on birthweight or growth. C & D) P21 female *Tor1a*^+/Δgag^ mice more frequently show gait defects than *Tor1a*^+/+^. * indicates a significant difference. Two-tailed Chi square test. E) Images of normal animal gait (upper panel), a female *Tor1a*^+/Δgag^ mouse with an out-turned hind paw (middle), and a female *Tor1a*^+/Δgag^ with abnormal hindlimb extension (lower).

We therefore considered whether *Tor1a*^+/Δgag^ acts with reduced penetrance *in utero* because different embryos experience different environments. For this we compared brain Lipin activity of *Tor1a*^+/+^ and *Tor1a*^+/Δgag^ embryos derived from dams fed a normal diet of 25% protein, versus dams fed an isocaloric low protein diet (8% protein). Maternal diet had no effect on brain Lipin activity of female or male *Tor1a*^+/+^ embryos, or male *Tor1a*^+/Δgag^ embryos (Figure 2D & E). However, female *Tor1a*^+/Δgag^ embryos from dams with low protein diet had significantly higher mean Lipin activity compared to those from dams on a normal diet. This amounted to double the number of female *Tor1a*^+/Δgag^ embryos with “pathological” brain Lipin activity (Figure 2E). We conclude that *in utero* environment modifies whether +/Δgag induces abnormal brain lipid metabolism.

### Dystonia-like symptoms in ~30% of female *Tor1a*^+/Δgag^ mice

+/Δgag induction of lipid metabolic defects is the first molecular event that occurs with reduced penetrance like DYT-*TOR1A* dystonia symptoms. We now examined whether it relates to disease. Although *in vitro* culturing of mouse cells poorly modeled the *in vivo* variability, we nevertheless examined human fibroblast lines from controls, a non-symptomatic *TOR1A*^+/Δgag^ individual, and three symptomatic *TOR1A*^+/Δgag^ patients (Supplemental Figure 4A). Baseline Lipin activity varied between lines without a clear correlation with genotype or symptomatic status. All *TOR1A*^+/Δgag^ lines had elevated *TOR1B* expression (Supplemental Figure 4B & C) that, when removed, led to similarly increased Lipin activity (Supplemental Figure 4D & E). Thus, it does not appear that *in vitro* cultured human cells can address the role of Lipin in disease.

We returned to *Tor1a*^+/Δgag^ mice. Previous profiling has suggested they lack overt dystonia-like symptoms (Tanabe et al., 2012; Ulug et al., 2011). However, studies focused on males. We now examined a sufficiently large cohort to detect behavioral defects of reduced penetrance and/or a role of sex. There was no growth difference between *Tor1a*^+/+^ and *Tor1a*^+/Δgag^ mice of either sex (Figure 3A & B), or the age they achieved a battery of sensory-motor reflexes, skills, and developmental milestones (Supplemental Figure 5). The diagnosis of DYT-*TOR1A* dystonia in humans is made from qualitative neurological assessment (Bressman et al., 2000). We reviewed movement, posture and gait of juvenile P21 *Tor1a*^+/Δgag^ mice, considering that DYT-*TOR1A* dystonia commonly presents as abnormal limb movements in childhood or adolescence (Bressman et al., 1994). Pairs of trained observers scored < 5% of *Tor1a*^+/+^ animals as abnormal. In contrast, this rose to 23% of *Tor1a*^+/Δgag^ (Figure 3C). All abnormal *Tor1a*^+/Δgag^ animals were female. This amounted to ~35% of females (Figure 3D), thus closely paralleling the sex and percentage of *Tor1a*^+/Δgag^ embryos with abnormal lipid metabolism. Video review identified two components to the motor dysfunction: (1) out-turned hind paws during ambulation (Figure 3E center panel) and, (2) abnormal hindlimb extension reminiscent of human dystonia (Figure 3E lower panel. Supplemental videos 1 - 2). Quantification of hind paw angles found a subset of female *Tor1a*^+/Δgag^ animals with wider mean paw angle than any female *Tor1a*^+/+^ (Supplemental Figure 6A & B). We compared the qualitative and quantitative analyses, which confirmed they identified the same *Tor1a*^+/Δgag^ females. No other *Tor1a*^+/Δgag^ females had abnormal values, suggesting that qualitative assessment detected all abnormal animals (Supplemental Figure 6B & C). We therefore conclude that overt neurological disease resembling dystonia occurs in *Tor1a*^+/Δgag^ mice with sex-skewed reduced penetrance.

### Lipin hyperactivity in development causes dystonia-like symptoms

We compared Lipin activity in brains of P21 female mice, but there was no difference between *Tor1a*^+/+^ and *Tor1a*^+/Δgag^ (Figure 4A). This is consistent with other reports that *Tor1a* mutations act most strongly in development (Tanabe et al., 2016), including that complete TorsinA loss no longer affects Lipin once mice reach 3-weeks (inset Figure 4B (Cascalho et al., 2020)). We assessed the penetrance that *Tor1a*^+/Δgag^ affects Lipin across development. 20-35% of female *Tor1a*^+/Δgag^ animals aged E13.5 to P14 had abnormally high Lipin activity, but the degree of hyperactivity peaked at E18.5 and reduced as animals matured (Figure 4B).

**Figure 4.**
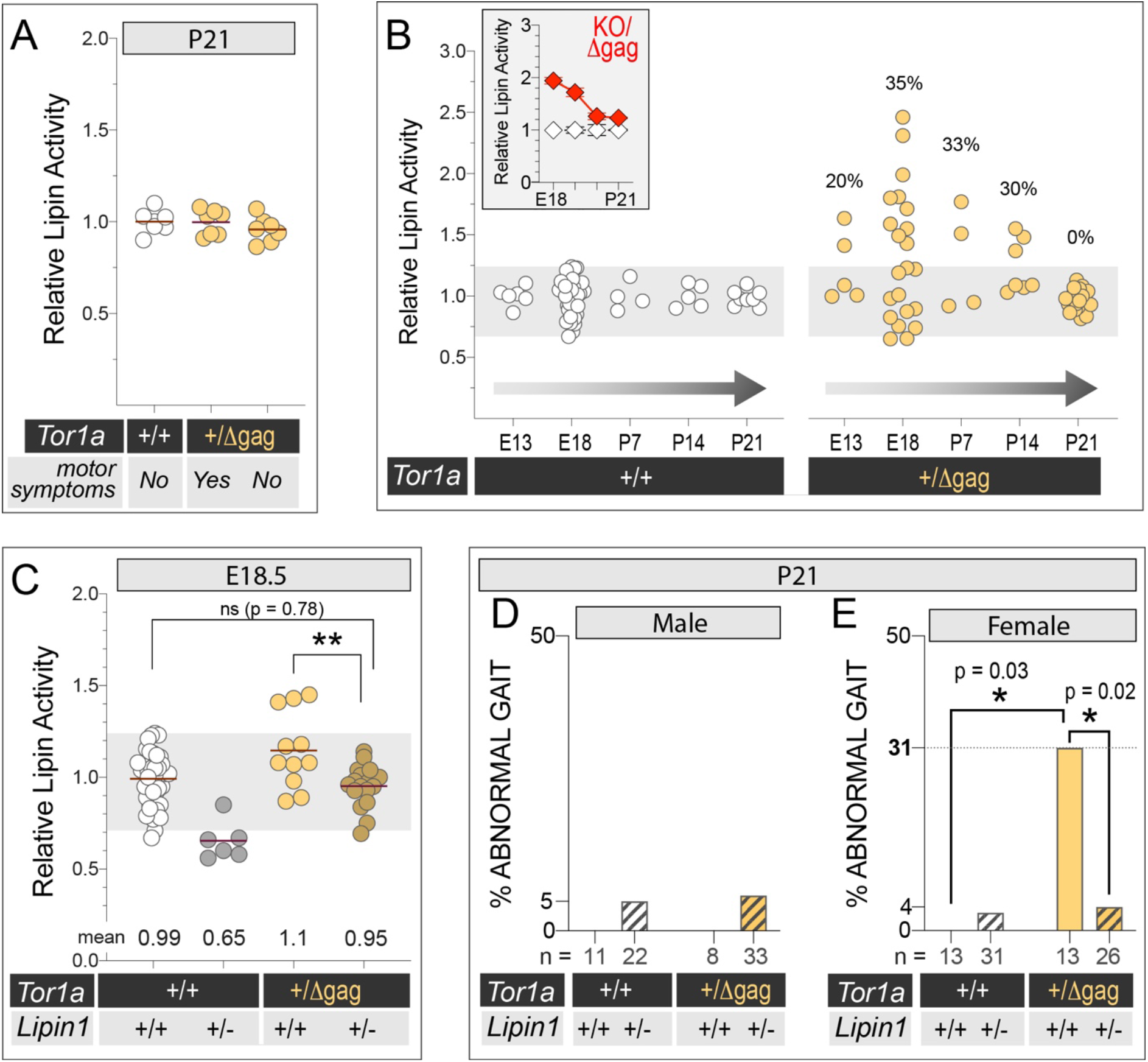
Lipin hyperactivity in development causes the movement disorder of P21 *Tor1a*^+/Δgag^ mice. A) Lipin activity in P21 female brain homogenates after behavioral testing shown in Figure 3. B) Lipin activity in female brain homogenates. The grey zone shows the range recorded for the *Tor1a*^+/+^ population, and numbers refer to the % of embryos with activity above this range. Inset shows Lipin activity of biallelic *Tor1a^KO^*^/Δgag^ mice, as first reported in (Cascalho et al., 2020). C) Lipin activity in E18.5 brain homogenates. Two-way ANOVA detects significant (p < 0.0001) effect of *Tor1a* and *Lipin1* genotypes. D & E) Frequency of gait defects in P21 mice. * indicates significant difference. Two-tailed Chi square test.

We tested whether this period of Lipin hyperactivity was responsible for the motor dysfunction in P21 mice. We interbred *Tor1a*^+/Δgag^ and *Lipin1*^+/−^ and collected embryos at E18.5. *Lipin1*^+/−^ brains had ~65% of normal Lipin activity. We compared *Tor1a*^+/Δgag^*:Lipin1*^+/+^ and *Tor1a*^+/Δgag^:*Lipin1*^+/−^. This showed that all *Tor1a*^+/Δgag^:*Lipin1*^+/−^ embryos had Lipin activity within the normal range (Figure 4C). We then examined motor function of a large cohort of P21 male and female mice. There was no difference between males (Figure 4D). We found significantly more female *Tor1a*^+/Δgag^:*Lipin1*^+/+^ with gait defects than *Tor1a*^+/+^:*Lipin1*^+/+^ (Figure 4E). This amounted to 31% of female *Tor1a*^+/Δgag^:*Lipin1*^+/+^ mice, tightly replicating the 35% of the first cohort. Gait defects were significantly affected by *Lipin1* genotype. Fewer female *Tor1a*^+/Δgag^:*Lipin1*^+/−^ mice had gait defects than *Tor1a*^+/Δgag^:*Lipin1*^+/+^, and in fact this was no different to the baseline recorded for *Tor1a*^+/+^:*Lipin1*^+/+^ controls (Figure 4E).

## Discussion

DYT*-TOR1A* dystonia has been recognized for more than twenty years. The disease has remained mysterious in terms of its molecular etiology, as well as its reduced penetrance; at least in part because +/Δgag did not appear to cause disease in animal models. Here we find previously undetected dystonia-like symptoms in *Tor1a*^+/Δgag^ mice. They develop in a subset of animals, thus mimicking the reduced penetrance of human DYT-*TOR1A* dystonia. Further, we show that a transient period of abnormal lipid metabolism during brain development defines which *Tor1a*^+/Δgag^ mice develop symptoms. These data therefore present a concrete example where an insult during development defines which genetically-susceptible individuals have disease in later life.

There is remarkable variability between inbred *Tor1a*^+/Δgag^ and *Tor1a*^+/−^ mice. Their inbred nature strongly suggests this is driven by variation in developmental processes and/ or *in utero* environment, rather than genetics. This is further supported by the fact that *Tor1a*^+/Δgag^ is haploinsufficient *in vitro*, as well as the impact of maternal nutrition. We do not solve the root cause why Lipin de-regulation only occurs in some embryos. The data best fit a model where *Tor1a*^+/Δgag^ embryos are compromised and unable to buffer Lipin activity if challenged by a second “hit”. This second hit preferentially affects female embryos, and thus female sex is a risk factor but does not invariably encode metabolic defects. The second hit is also more likely under nutrient stress. Indeed, the general principle of a second hit is illustrated by *Tor1a*^+/Δgag^ fibroblasts maintaining normal Lipin activity until additionally compromised by *TOR1B*/*Tor1b* depletion.

It is clear that *Tor1b* is not the second hit driver of Lipin hyperactivity during *Tor1a*^+/Δgag^ brain development. Moreover, genome-wide RNAseq shows remarkably stable gene transcription between affected and unaffected animals. This suggests the second hit acts on Lipin by a post-translational mechanism; in fact consistent with the complex post-translational regulation of Lipin enzymes (Eaton et al., 2013; Harris et al., 2007; Hsieh et al., 2015; Li et al., 2018; Liu and Gerace, 2009). Additional work is needed to dissect which pathways regulate Lipin during brain development, how they are affected by developmental events and stresses, and their relationship to sex. However, it is intriguing to consider that low protein diet increases the number of affected *Tor1a*^+/Δgag^ embryos. Low protein diet also suppresses mTOR signaling which, in turn, dis-inhibits mTORC1 regulation of Lipin1 (Lamming et al., 2015; Peterson et al., 2011).

It is surprising that +/Δgag preferentially causes a movement disorder in female mice, but similarly affects male and female humans (Martino et al., 2013). This difference arises *in utero* suggesting it relates to pregnancy differences between species; potentially the multiple pregnancies of mice where male and female embryos differentially compete for resources (Bale, 2016). Nevertheless, the sex skewing of the movement disorder in mice is intriguing because many forms of dystonia preferentially occur in female children and women. GCH1 mutations are two to four times more likely to cause Dopa-Responsive Dystonia in female children (Furukawa et al., 1998; Wijemanne and Jankovic, 2015). This form of dystonia also arises before puberty, thus pointing to female sex rather than societal factors as a risk factor for dystonia.

These data caution against the practice of focusing on male rodents. This remains prevalent, including that in 2016 and 2017, 75% of drug studies only studied male mouse behavior (Hughes, 2019). This bias is detrimental for women’s health (Beery and Zucker, 2011). Additionally, here we show a surprising example where sex more strongly influences pathophysiology in mice than men. It took a comprehensive analysis of male and female *Tor1a*^+/Δgag^ mice to identify that females are an exceptionally accurate animal model for a disease that had been considered as challenging to mimic. This finding already showed that Lipin hyperactivity has a key role in disease pathogenesis. Moreover, it provides dystonia research with a powerful tool to further define disease etiology and thus drive novel therapeutic interventions.

## Acknowledgments

This work was only possible by the support of the Foundation for Dystonia Research. We thank the Dystonia Medical Research Foundation for an award. A.C. and J.F. have FWO Sb fellowships (1S58816N and 1S54119N). We thank the VIB Nucleomics Facility for RNAseq. We make a major acknowledgement to Lise Hennebel, and to collaborations with Christine Klein and Philip Seibler (Institute of Neurogenetics, University of Luebeck) for fibroblast lines, as well as Antonio Pisani (University of Rome) for many helpful insights.

## Author contributions

A.C., J.F. and R.E.G. were involved in project conceptualization, data collection and analysis. A.C. and R.E.G. supervised experiments. J.F. and R.E.G. prepared the manuscript. S.R., N.V. and S.F.G. performed experiments.

## Methods

### Mouse lines, husbandry, and tissue collection

The *Tor1a* Δgag and KO alleles are previously described (Goodchild et al., 2005) and have been backcrossed more than 20-times onto the C57BL/6J background. The *Lipin1* KO allele was acquired from Jackson Mice (Peterfy et al., 2001) and was crossed at least 5 times onto the C57BL/6J background. Biochemical and mRNA measurements were performed on tissues collected from animals derived from crossing *Tor1a*^+/Δgag^ males with C57BL/6J females. Breeding females were checked daily for mating based on the presence of a vaginal plug. They were then single housed with *ad lib* access to water and Ssniff® M-Z pellets or Ssniff®EF R/M Protein Deficient pellets for nutritional challenge experiments (Spezialdiäten GmbH). TransnetYX performed genotyping for *Tor1a, Lipin1* and the Y-chromosome, as well as the SNP analysis of genetic background

The day a vaginal plug was detected is considered as E0.5. Embryos were collected from pregnant females after they were euthanized by cervical dislocation. Days of post-natal development were defined after assigning the day of birth as P0. Postnatal animals were permanently identified by tattooing. Tissues used in biochemistry and mRNA analyses were collected from embryos and animals aged P0-P14 after they were euthanized by decapitation, or by cervical dislocation for animals older than P14. These tissues were snap frozen in liquid nitrogen and stored at −80°C until use. Unless stated otherwise, all brains were separated into left and right hemispheres before freezing to allow two analyses per animal.

All mice were housed in the KU Leuven animal facility that is fully compliant with European policy on the use of Laboratory Animals. All animal procedures were approved by the Institutional Animal Care and Research Advisory Committee of the KU Leuven (ECD P060/2017) and performed in accordance with the Animal Welfare Committee guidelines of KU Leuven, Belgium.

### Cell Lines and culturing

MEFs were produced from E13.5 litters and immortalized with the SV40 large tumor-antigen as previously described (Cascalho et al., 2020). Human fibroblasts were acquired from the NINDS repository (https://nindsgenetics.org), or were previously described (Cascalho et al., 2020). Lines were cultured in Dulbecco’s Modified Eagle Media (DMEM; Thermo Fisher Scientific) containing 10% Fetal Bovine Serum (characterized FBS; Hyclone). shRNA plasmids were acquired from Sigma: Mouse *Tor1b* (TRCN0000106485), Human *TOR1B* (TRCN0000159398) or control (Empty pLKO.1) and introduced into cell lines as previously described (Cascalho et al., 2020). Electroporated cells were plated into flasks and cultured for 72 hours before harvesting by trypsinization. The cell suspensions were separated into aliquots, washed, and pelleted. One aliquot was used for mRNA measurements and one for biochemical assays.

### Lipid enzyme assays

Snap frozen brain hemispheres or cell pellets were homogenized on ice in “PAP enzyme buffer” (50mM Tris HCl (pH 7.5), 0.25M sucrose, 10mM 2-mercaptoethanol, 1x PhosSTOP phosphatase inhibitor cocktail (Roche) and 1x EDTA free protease inhibitor cocktail (Sigma)). Debris was removed by centrifugation for 10 min at 1,000 x g at 4 °C, and supernatant was transferred to a fresh Eppendorf tube. The protein concentration of each homogenate was measured using a Bradford assay according to the manufacturer’s instructions (Bio-Rad).

PAP activity was measured according to standard protocols (Cascalho et al., 2020; Dubots et al., 2014; Sembongi et al., 2013) based on the formation of fluorescent DAG from NBD-PA (Avanti® Polar lipids, Inc). In brief, assays used 60μg of total protein diluted in PAP enzyme buffer to achieve final concentrations of 2mM NBD-PA, and either 0.5mM MgCl_2_ or 1mM EDTA. These reactions were incubated for 30 min at 30°C. Lipids were extracted and separated by Thin Layer Chromatography, and fluorescence was imaged with an ImageQuant LAS 4000 device (Green-RGB, 460nm/534nm). ImageJ was used to quantify levels of PA and DAG fluorescence. Total PAP activity was calculated as the amount of NBD-DAG relative to all NBD-labeled lipids (PA+DAG) in the MgCl_2_ condition. LPP PAP activity was calculated as the amount of NBD-DAG relative to all NBD-labeled lipids (PA+DAG) in the EDTA-containing condition. Lipin activity was calculated as the difference (Supplemental Figure 1A). DAG Kinase activity was measured based on the production of fluorescent PA from NBD-DAG (Avanti® Polar lipids, Inc) in the same buffer condition used to measure total PAP activity. DAG Kinase activity was calculated as the fraction of NBD-PA relative to the total NBD-labeled lipids (PA+DAG).

### Protein and Transcriptional Analyses

Western blotting was performed with brain homogenates prepared for PAP enzyme assays. 30μg of protein was subject to standard SDS-PAGE, transferred to a PVDF membrane, and blocked (5% milk, 0.2% Tween, 1x PBS). Rabbit anti-torsinA and anti-torsinB are custom generated antibodies. We previously defined conditions where they specifically recognize their respective antigens (Jungwirth et al., 2010). Membranes were incubated overnight at 4°C with primary antibodies in blocking buffer, washed in buffer (0.2% Tween, 1x PBS), and incubated with horseradish peroxidase conjugated anti-rabbit antibody diluted in blocking buffer (Jackson Immunoreagents: 715-035-150). They then received a final series of washing steps, followed by incubation with West Pico Plus chemiluminescent reagent (ThermoFisher Scientific). Chemiluminescence was detected with an ImageQuant LAS 4000 device.

RNA was extracted from cells and tissues after preservation in RNAlater (Qiagen). In all cases, samples were homogenized and RNA was purified using a QIAshredder and RNeasy Qiagen Mini Kit according to the manufacturer’s instructions (Qiagen). The RNAseq profiling was previously described (Cascalho et al., 2020). mRNA expression in cell lines was analysed after producing cDNA with SuperScript IV Reverse Transcriptase (ThermoFisher Scientific) and 50μM random hexamer priming. qPCR was performed with 500ng of cDNA and the SensiFast_™_ SYBR_®_ No-ROX kit (Bioline) and a Lightcycler® 480/1536 (Roche) under the SYBRGreen standard run protocol. All qPCR runs were performed in duplicate. Primers and conditions have been previously described (Cascalho et al., 2020).

### Behavioral assays

The acquisition of behavioral skills was examined using established protocols (Feather-Schussler and Ferguson, 2016). Animal gait was assessed in individual animals placed in a fresh cage facing away from the observer over at least 2 minutes. A score of NORMAL was assigned to animals that moved with both hindlimbs participating evenly, body weight supported on all four paws, and if the abdomen did not touch the ground. An animal was scored as ABNORMAL if tremor or limping were observed, if the pelvis was lowered, feet or limbs pointed away from the body during locomotion (“duck feet”), if an animal had difficulty moving forward, or if it dragged its abdomen. These analyses were performed pre-weaning on all pups present in a nest. The first analysis of P21 *Tor1a*^+/+^ and *Tor1a*^+/Δgag^ gait was performed on pups produced by breeding *Tor1a*^+/Δgag^ males with C57BL6/J dams. The postnatal development of *Tor1a*^+/+^ and *Tor1a*^+/Δgag^ animals was studied using pups produced by crossing *Tor1a^flox/flox^*:*Lipin1*^+/−^ dams with *Tor1a*^+/Δgag^:*Lipin1*^+/−^:*Nestin-Cre* males, as described in (Cascalho et al., 2020). No *Nestin-Cre* transgene carrying offspring or *Lipin1*^+/−^ offspring were included in these analyses. The cohort of P21 *Tor1a*^+/+^:*Lipin1*^+/+^, *Tor1a*^+/+^:*Lipin1*^+/−^, *Tor1a*^+/Δgag^:*Lipin1*^+/+^ and *Tor1a*^+/Δgag^:*Lipin1*^+/−^ mice were also derived from the *Tor1a^flox/flox^*:*Lipin1*^+/−^ x *Tor1a*^+/Δgag^: *Lipin1*^+/−^:*Nestin-Cre* crosses and again no data from *Nestin-Cre* transgene offspring were included in analyses.

### Statistical analysis

N values are given in figures or legends and reflect the number of individual animals unless specifically stated otherwise. No measurements were excluded from any analysis. All analyses were performed blind to genotype and sex. Unless stated otherwise, statistical analyses were performed using Prism 8.3 (GraphPad Software). Tukey’s multiple comparison test was used for all post hoc comparisons. *, **, ***, and **** indicate significant differences between groups at p < 0.05, 0.01, 0.001, and 0.0001 respectively. Tests are stated in the figure legends.

---

**Supplemental Figure 1.**
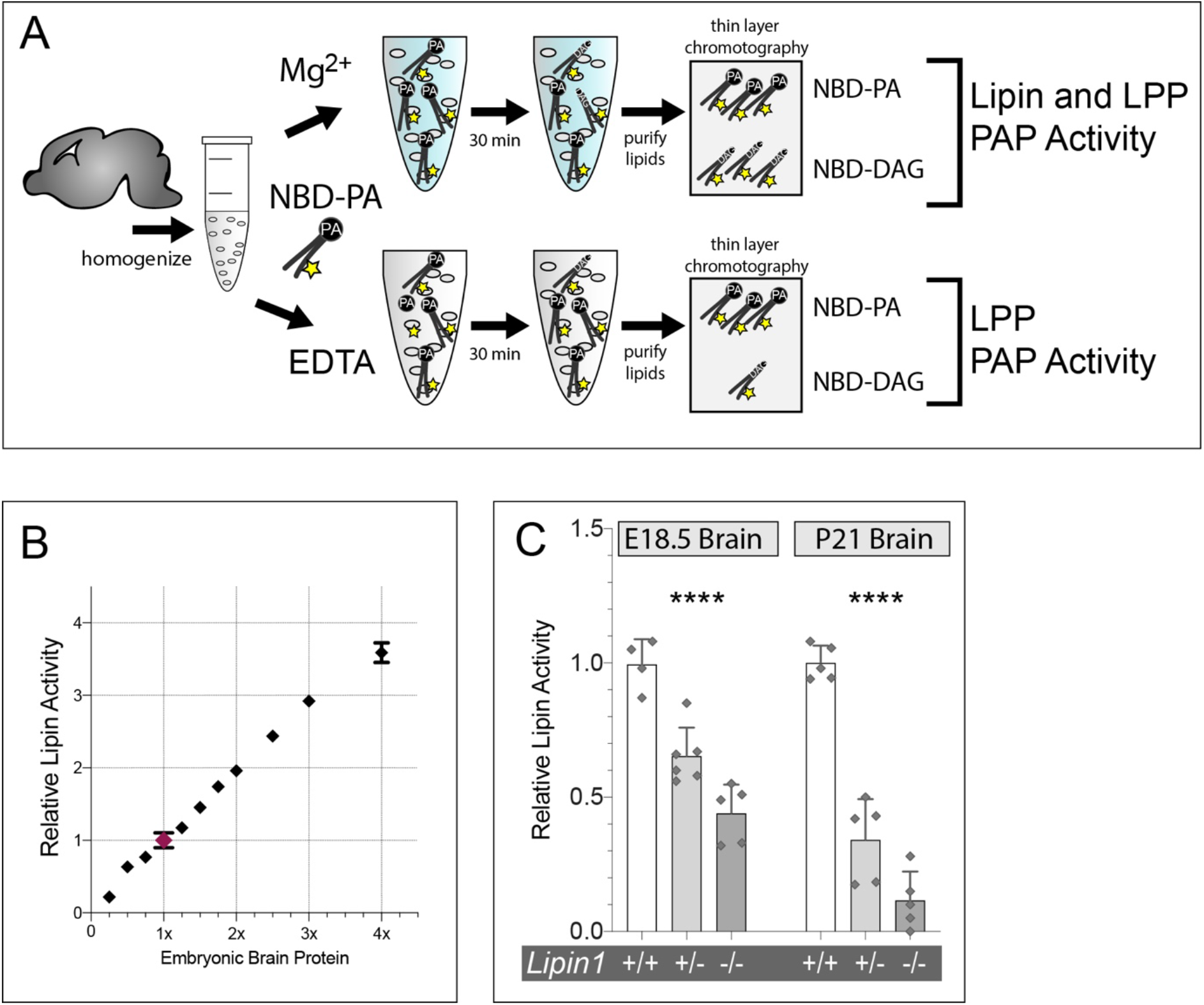
Quantitation of Lipin activity. A) Schematic of the PAP enzyme activity. Lipin activity is calculated from DAG production in the presence of magnesium, minus DAG production in the EDTA condition. B) Linearity of the Lipin assay. Lipin values are expressed relative to the activity found in “1x” (60μg) brain protein that is used as standard in all assays. There is a linear relationship between Lipin activity measurements and brain protein ranging from 0.25x (15μg) to 3.0x (180μg) of this standard; above this the curve flattened. All *Tor1a*^+/+^, *Tor1a*^+/Δgag^, and *Tor1a*^+/−^ Lipin activity values lie within this linear range. C) *Lipin1* gene deletion reduces Lipin activity in brain homogenates from embryos and post-natal animals. Data is presented relative to *Lipin1*^+/+^ at each age. Points show values from individual animals, bars show mean +/− SD. **** indicates that Two-Way ANOVA detects a significant effect of *Lipin1* genotype (P<0.0001). Note that embryonic brain expresses *Lipin1* and *Lipin2*.

**Supplemental Figure 2.**
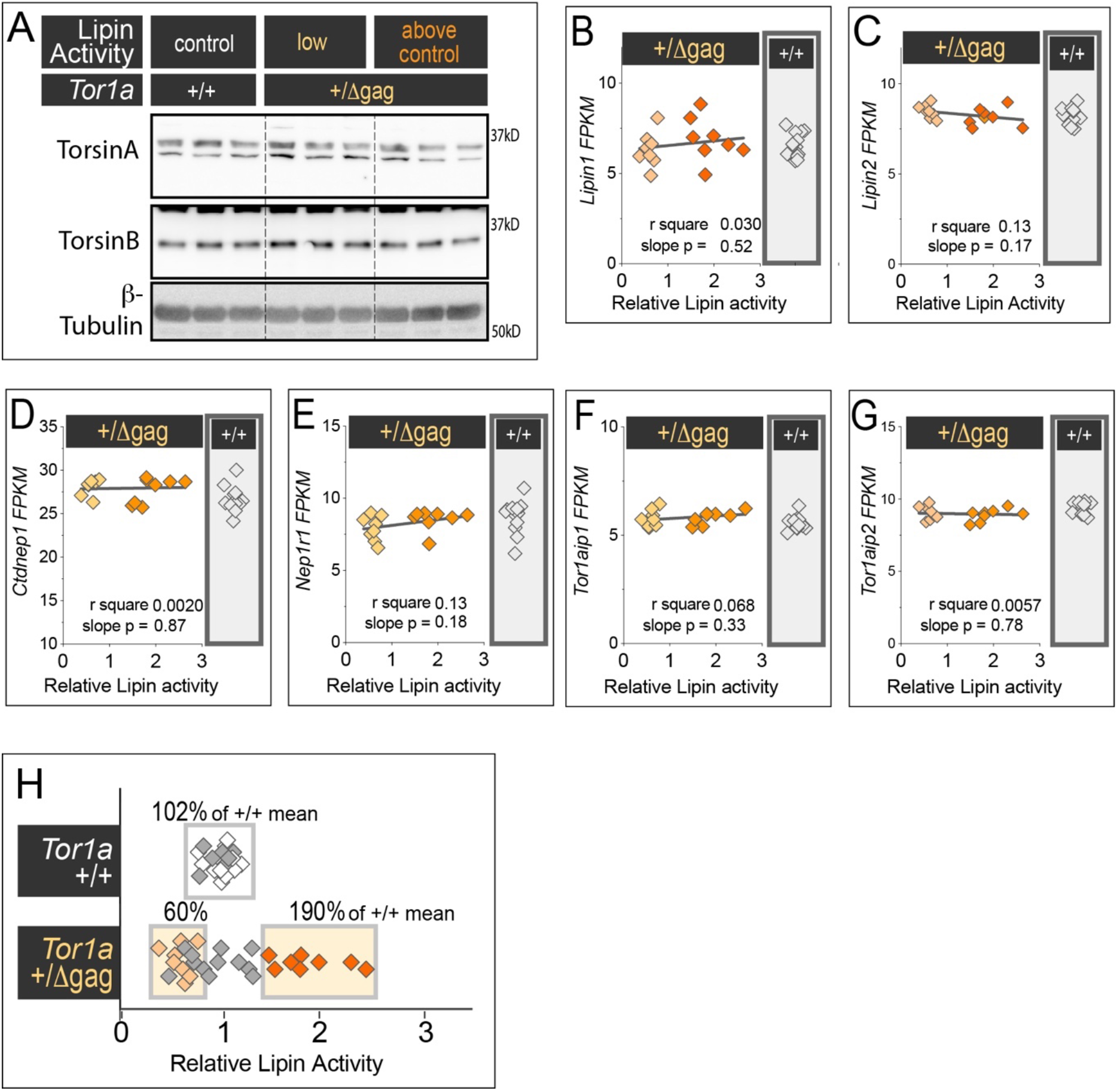
Varied gene expression does not account for the variation in Lipin activity between *Tor1a*^+/Δgag^ brains. A) Western blotting finds similar TorsinA and TorsinB levels in the brains of E18.5 *Tor1a*^+/+^ and *Tor1a*^+/Δgag^ embryos, and between *Tor1a*^+/Δgag^ embryos with different levels of brain Lipin PAP activity. Each lane shows immunoreactivity of one embryo, with three embryos assessed per group. We previously validated that anti-TorsinA and anti-TorsinB immunoreactivity reflect TorsinA and TorsinB protein levels (Jungwirth et al., 2010). B – G) Lipin activity plotted against the FPKM of each gene for brains from individual E18.5 *Tor1a*^+/Δgag^ embryos. The panel on the right shows data for individual *Tor1a*^+/+^ animals. H) Brain Lipin activity in E18.5 embryos selected for RNAseq analysis. Boxes delineate the three groups, % values show the mean Lipin activity for each group relative to the *Tor1a*^+/+^ mean.

**Supplemental Figure 3.**
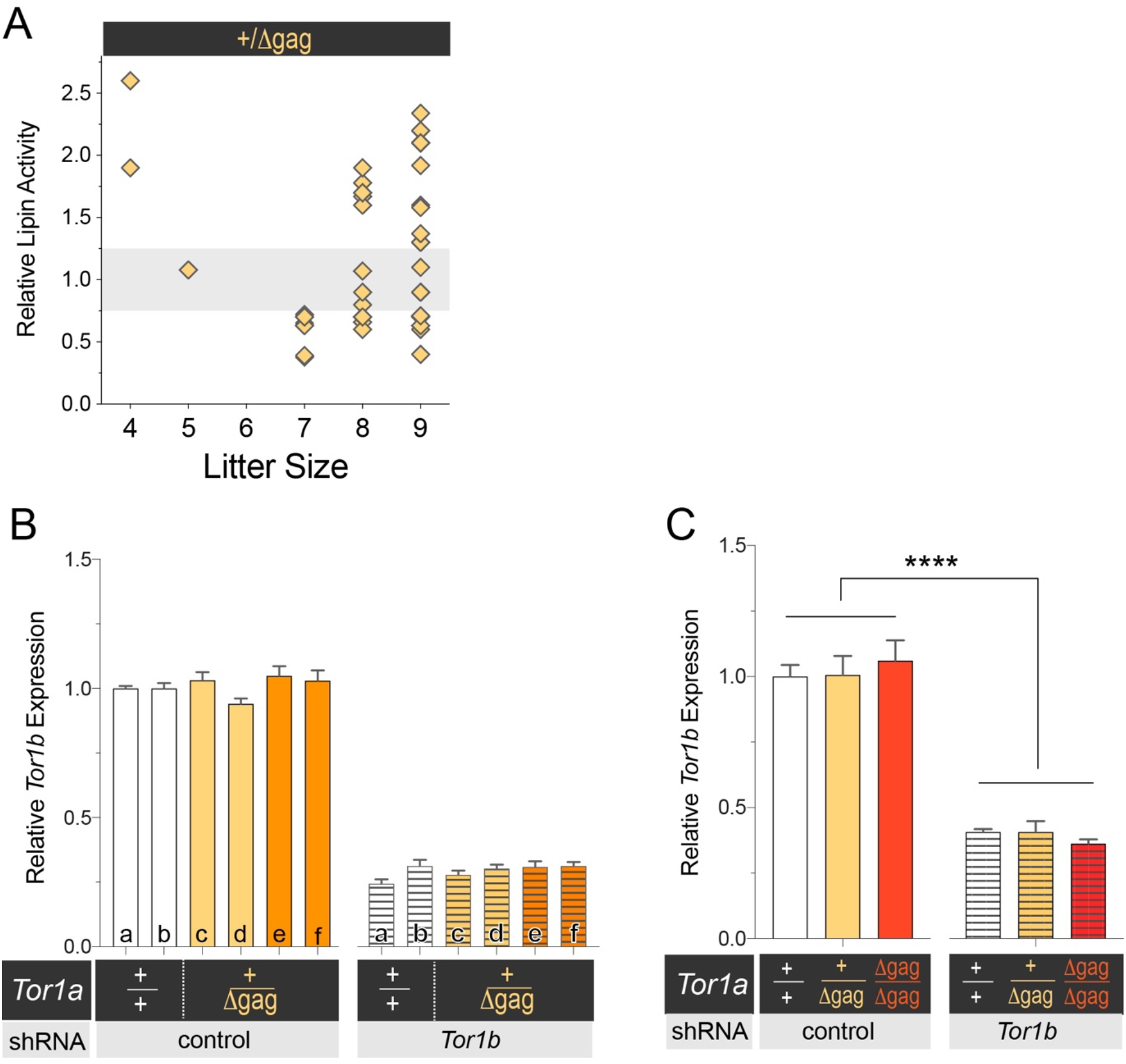
*Tor1a*^+/Δgag^ affects lipid metabolism with reduced penetrance *in vivo* but not *in vitro*. A) *Tor1a*^+/Δgag^ embryos with Lipin hyperactivity are present in small and large litters. Points show brain Lipin activity of individual E18.5 *Tor1a*^+/Δgag^ embryos plotted by the number of embryos per litter. Shaded area highlights the range of brain Lipin activity from the *Tor1a*^+/+^ embryo population (not shown). B) *Tor1b* mRNA expression in MEF lines derived from *Tor1a*^+/+^ and *Tor1a*^+/Δgag^ embryos, and electroporated with empty vector or *Tor1b* shRNA vector. Data is presented relative to the mean of *Tor1a*^+/+^ with empty vector. Bars show mean +/− SEM of duplicate electroporations per line. Letters refer to individual embryos shown in Figure 2A C) *Tor1b* mRNA expression in MEF lines derived from *Tor1a*^+/+^, *Tor1a*^+/Δgag^ and *Tor1a*^Δgag/Δgag^ embryos, and electroporated with empty vector or *Tor1b* shRNA vector. Data is presented relative to the mean of *Tor1a*^+/+^ with empty vector. Two-way ANOVA detects a significant effect of *Tor1b* knockdown (****, p < 0.0001) and no significant effect of *Tor1a* genotype.

**Supplemental Figure 4.**
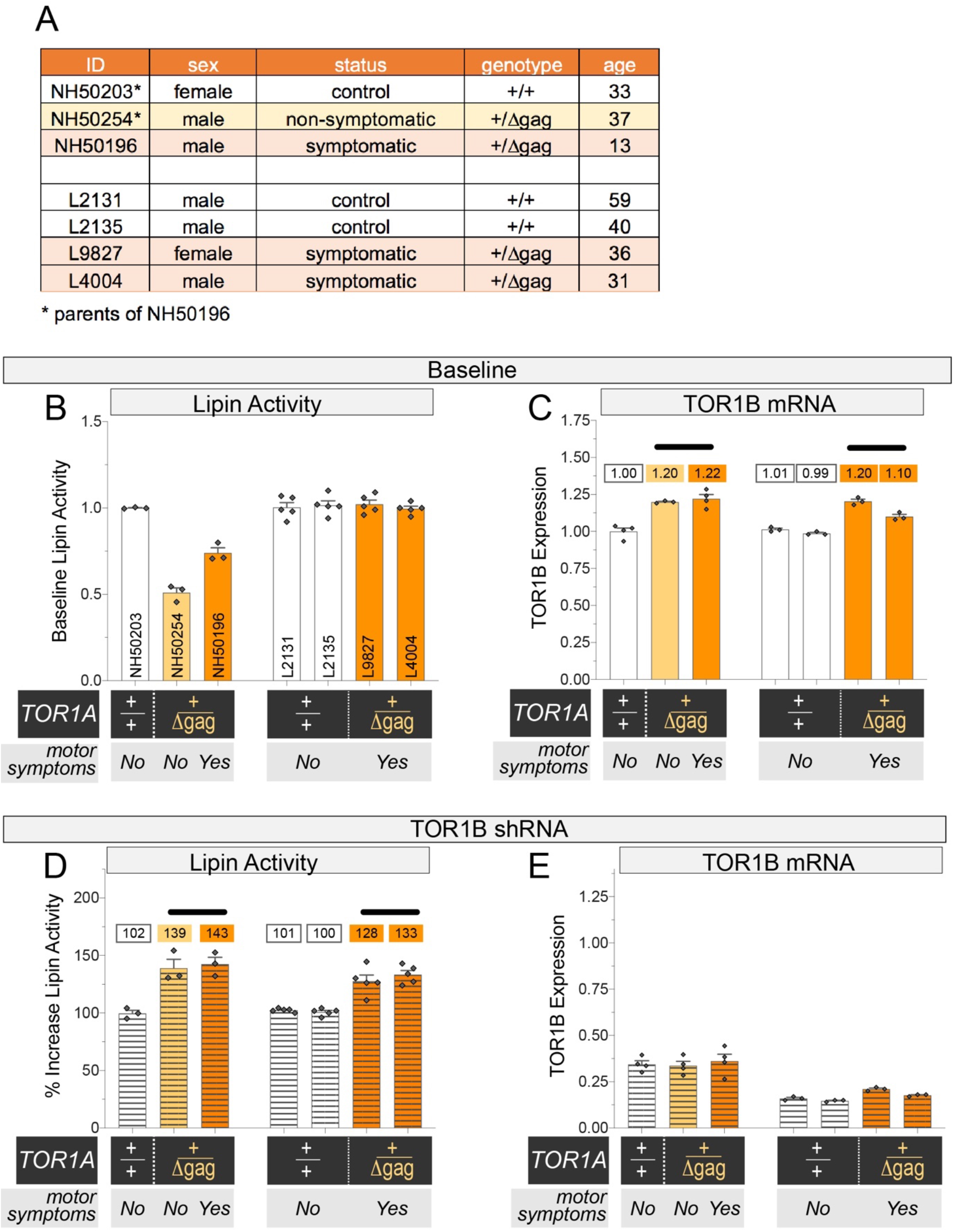
Human fibroblast line *TOR1B* expression and Lipin activity. A) Human fibroblast cell line characteristics. B) Lipin activity at baseline in homogenates prepared from human fibroblast lines after electroporation with empty vector. Bars show mean +/− SEM, and points show values from each electroporation. Vertical numbers refer to the ID code of each line, and the same series is repeated in figure panels C-E. C) *TOR1B* expression measured by qRT-PCR after electroporation with empty vector. Numbers show the mean relative *TOR1B* expression for each line. Unpaired Two-tailed T-Test detects a significant difference between *TOR1A*^+/+^ and *TOR1A*^+/Δgag^ lines (p = 0.003). D) Percentage change in Lipin activity between human fibroblast cell lines electroporated with *TOR1B* shRNA vector, and baseline *TOR1B* expression measured from the empty vector condition. Unpaired Two-tailed T-Test detects a significant difference between *TOR1A*^+/+^ and *TOR1A*^+/Δgag^ lines (p = 0.0003). E), as (C), but following electroporation with *TOR1B* shRNA vector. There is no difference in *TOR1B* expression between *TOR1A*^+/+^ and *TOR1A*^+/Δgag^ lines.

**Supplemental Figure 5.**
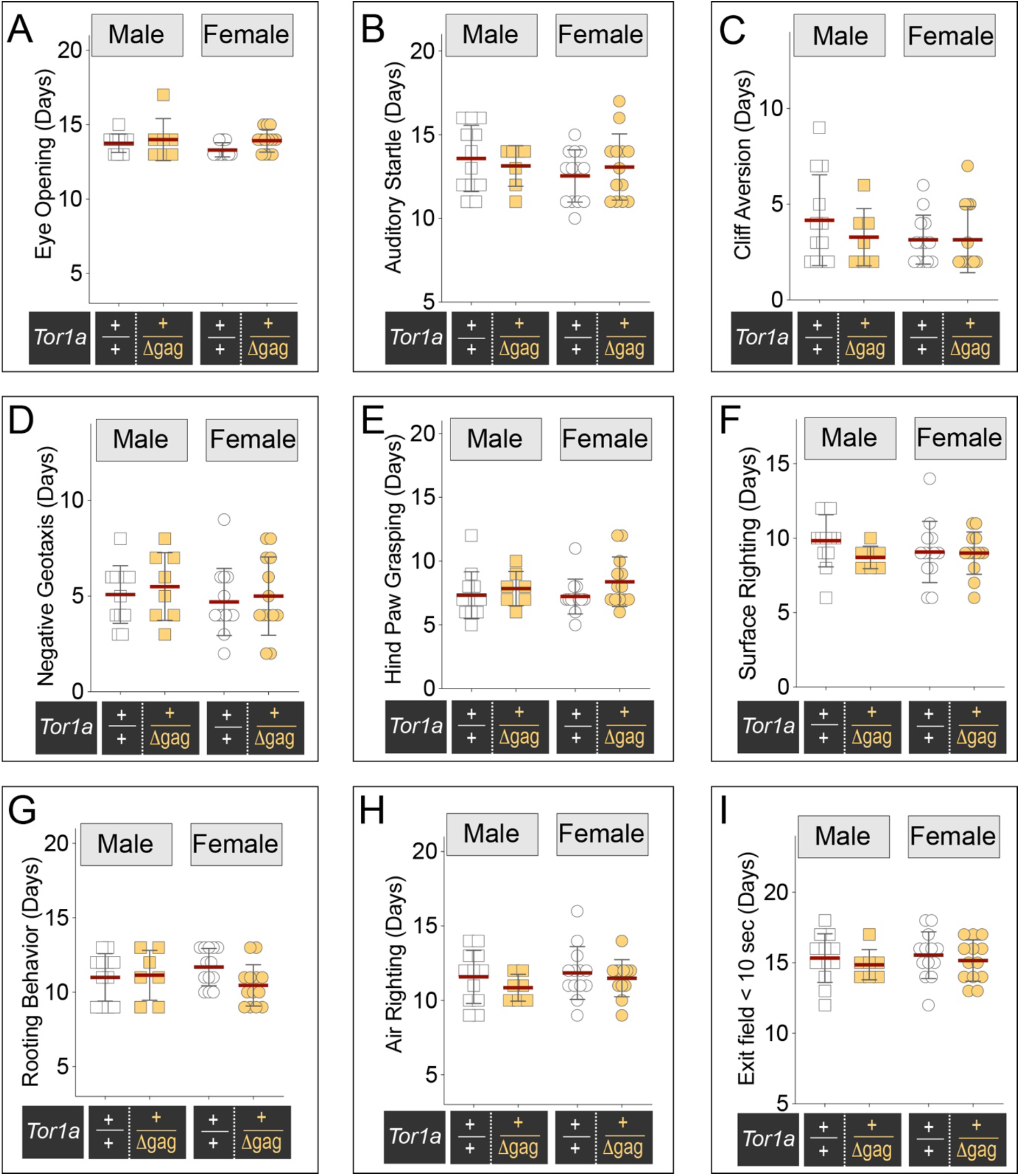
*Tor1a*^+/Δgag^ animals progress normally to developmental milestones. Points show the age that individual animals achieve a reflex, behavior or developmental event. Bars show mean +/− SD. A) Age of animal (postnatal days) when both eyes are open. Two-way ANOVA finds no effect of *Tor1a* genotype (F (1, 41) = 3.1), sex (F (1, 41) = 1.1), or an interaction between *Tor1a* genotype and sex (F (1, 41) = 0.56). B) Age of animal (postnatal days) when it first reacts to an auditory stimulus. Two-way ANOVA finds no effect of *Tor1a* genotype (F (1, 41) = 0.009), sex (F (1, 41) = 1.4), or an interaction between *Tor1a* genotype and sex (F (1, 41) = 0.81). C) Age of animal (postnatal days) when it first successfully crawls away from an elevated edge. Two-way ANOVA finds no effect of *Tor1a* genotype (F (1, 41) = 0.64), sex (F (1, 41) = 1.1), or an interaction between *Tor1a* genotype and sex (F (1, 41) = 0.64). D) Age of animal (postnatal days) when it is first able to reverse to face upward after being placed facing downward on a 45° slope. Two-way ANOVA finds no effect of *Tor1a* genotype (F (1, 41) = 0.01), sex (F (1, 41) = 0.69), or an interaction between *Tor1a* genotype and sex (F (1, 41) = 0.46). E) Age of animal (postnatal days) when it first shows a grasping reflex with hind paws. Two-way ANOVA finds no effect of *Tor1a* genotype (F (1, 41) = 2.6), sex (F (1, 41) = 0.17), or an interaction between *Tor1a* genotype and sex (F (1, 41) = 0.37). F) Age of animal (postnatal days) when it is first able to flip (< 2 sec) upright from a supine position. Two-way ANOVA finds no effect of *Tor1a* genotype (F (1, 41) = 1.4), sex (F (1, 41) = 0.2), or an interaction between *Tor1a* genotype and sex (F (1, 41) = 1.0). G) Age of animal (postnatal days) when it first shows a rooting reflex by moving its head towards a light tactile stimulus. Two-way ANOVA finds no effect of *Tor1a* genotype (F (1, 41) = 1.4), sex (F (1, 41) = 0.00), or an interaction between *Tor1a* genotype and sex (F (1, 41) = 2.3). H) Age of animal (postnatal days) when it first achieves a mid-air righting reflex by landing on all four paws when dropped from 10 cm height onto a soft surface. Two-way ANOVA finds no effect of *Tor1a* genotype (F (1, 40) = 1.2), sex (F (1, 40) = 0.89), or an interaction between *Tor1a* genotype and sex (F (1, 40) = 0.16). I) Age of animal (postnatal days) when it first ambulates and explores rapidly enough to exit a 10cm diameter zone in under 10 seconds. Two-way ANOVA finds no effect of *Tor1a* genotype (F (1, 41) = 0.81), sex (F (1, 41) = 0.28), or an interaction between *Tor1a* genotype and sex (F (1, 41) = 0.009).

**Supplemental Figure 6.**
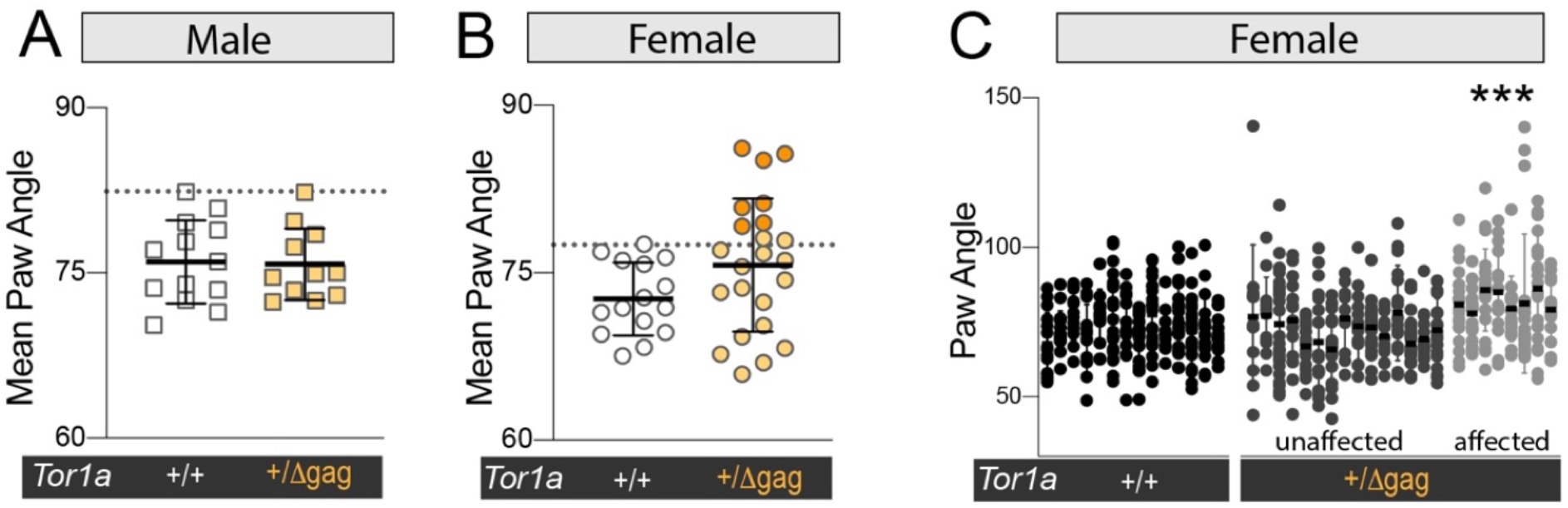
Quantification of hind-paw placement in P21 *Tor1a*^+/+^ and *Tor1a*^+/Δgag^ mice. A & B) Quantification of hind-paw angle during walking. Points show the mean hind paw angle of individual P21 mice, with at least 5 measurements per animal. Dotted lines show the widest mean angle recorded from *Tor1a*^+/+^ animals. (A) All male *Tor1a*^+/Δgag^ are within the range of male *Tor1a*^+/+^. (B) A subset of female *Tor1a*^+/Δgag^ have wider mean hind-paw angles than any *Tor1a*^+/+^ female. Orange points indicate which animals were scored as qualitatively abnormal. C) Hind-paw angles from individual *Tor1a*^+/+^ and *Tor1a*^+/Δgag^ females. Values from individual mice are arranged vertically, and the mean from each animal is highlighted by a dark bar. “Affected” and “unaffected” refer to the qualitative scoring of animals. *** indicates significance difference between *Tor1a*^+/Δgag^ that were qualitatively scored as abnormal, and *Tor1a*^+/+^ and *Tor1a*^+/Δgag^ that appeared normal. One-way ANOVA.

## References

Bale, T.L. (2016). The placenta and neurodevelopment: sex differences in prenatal vulnerability. Dialogues Clin Neurosci 18, 459–464.

Beery, A.K., and Zucker, I. (2011). Sex bias in neuroscience and biomedical research. Neurosci Biobehav Rev 35, 565–572.

Bressman, S.B., de Leon, D., Kramer, P.L., Ozelius, L.J., Brin, M.F., Greene, P.E., Fahn, S., Breakefield, X.O., and Risch, N.J. (1994). Dystonia in Ashkenazi Jews: clinical characterization of a founder mutation. Ann Neurol 36, 771–777.

Bressman, S.B., Sabatti, C., Raymond, D., de Leon, D., Klein, C., Kramer, P.L., Brin, M.F., Fahn, S., Breakefield, X., Ozelius, L.J., et al. (2000). The DYT1 phenotype and guidelines for diagnostic testing. Neurology 54, 1746–1752.

Cascalho, A., Foroozandeh, J., Hennebel, L., Swerts, J., Klein, C., Rous, S., Dominguez Gonzalez, B., Pisani, A., Meringolo, M., Gallego, S., et al. (2020). Excess Lipin enzyme activity contributes to TOR1A recessive disease and DYT-TOR1A dystonia. Brain. *In Press*.

Cooper, D.N., Krawczak, M., Polychronakos, C., Tyler-Smith, C., and Kehrer-Sawatzki, H. (2013). Where genotype is not predictive of phenotype: towards an understanding of the molecular basis of reduced penetrance in human inherited disease. Hum Genet 132, 1077–1130.

Craddock, C.P., Adams, N., Bryant, F.M., Kurup, S., and Eastmond, P.J. (2015). PHOSPHATIDIC ACID PHOSPHOHYDROLASE Regulates Phosphatidylcholine Biosynthesis in Arabidopsis by Phosphatidic Acid-Mediated Activation of CTP:PHOSPHOCHOLINE CYTIDYLYLTRANSFERASE Activity. Plant Cell 27, 1251–1264.

Demircioglu, F.E., Sosa, B.A., Ingram, J., Ploegh, H.L., and Schwartz, T.U. (2016). Structures of TorsinA and its disease-mutant complexed with an activator reveal the molecular basis for primary dystonia. Elife 5.

Dubots, E., Cottier, S., Peli-Gulli, M.P., Jaquenoud, M., Bontron, S., Schneiter, R., and De Virgilio, C. (2014). TORC1 regulates Pah1 phosphatidate phosphatase activity via the Nem1/Spo7 protein phosphatase complex. PLoS One 9, e104194.

Eaton, J.M., Mullins, G.R., Brindley, D.N., and Harris, T.E. (2013). Phosphorylation of lipin 1 and charge on the phosphatidic acid head group control its phosphatidic acid phosphatase activity and membrane association. J Biol Chem 288, 9933–9945.

Feather-Schussler, D.N., and Ferguson, T.S. (2016). A Battery of Motor Tests in a Neonatal Mouse Model of Cerebral Palsy. J Vis Exp.

Frederic, M.Y., Clot, F., Blanchard, A., Dhaenens, C.M., Lesca, G., Cif, L., Durr, A., Vidailhet, M., Sablonniere, B., Calender, A., et al. (2009). The p.Asp216His TOR1A allele effect is not found in the French population. Mov Disord 24, 919–921.

Furukawa, Y., Lang, A.E., Trugman, J.M., Bird, T.D., Hunter, A., Sadeh, M., Tagawa, T., St George-Hyslop, P.H., Guttman, M., Morris, L.W., et al. (1998). Gender-related penetrance and de novo GTP-cyclohydrolase I gene mutations in dopa-responsive dystonia. Neurology 50, 1015–1020.

Gluckman, P.D., Hanson, M.A., and Mitchell, M.D. (2010). Developmental origins of health and disease: reducing the burden of chronic disease in the next generation. Genome Med 2, 14.

Goodchild, R.E., Kim, C.E., and Dauer, W.T. (2005). Loss of the Dystonia-Associated Protein TorsinA Selectively Disrupts the Neuronal Nuclear Envelope. Neuron 48, 923–932.

Grillet, M., Dominguez Gonzalez, B., Sicart, A., Pottler, M., Cascalho, A., Billion, K., Hernandez Diaz, S., Swerts, J., Naismith, T.V., Gounko, N.V., et al. (2016). Torsins Are Essential Regulators of Cellular Lipid Metabolism. Dev Cell 38, 235–247.

Harris, T.E., Huffman, T.A., Chi, A., Shabanowitz, J., Hunt, D.F., Kumar, A., and Lawrence, J.C., Jr. (2007). Insulin controls subcellular localization and multisite phosphorylation of the phosphatidic acid phosphatase, lipin 1. J Biol Chem 282, 277–286.

Hsieh, L.S., Su, W.M., Han, G.S., and Carman, G.M. (2015). Phosphorylation regulates the ubiquitin-independent degradation of yeast Pah1 phosphatidate phosphatase by the 20S proteasome. J Biol Chem 290, 11467–11478.

Huffman, T.A., Mothe-Satney, I., and Lawrence, J.C., Jr. (2002). Insulin-stimulated phosphorylation of lipin mediated by the mammalian target of rapamycin. Proc Natl Acad Sci U S A 99, 1047–1052.

Hughes, R.N. (2019). Sex still matters: has the prevalence of male-only studies of drug effects on rodent behaviour changed during the past decade? Behav Pharmacol 30, 95–99.

Jungwirth, M., Dear, M.L., Brown, P., Holbrook, K., and Goodchild, R. (2010). Relative tissue expression of homologous torsinB correlates with the neuronal specific importance of DYT1 dystonia-associated torsinA. Hum Mol Genet 19, 888–900.

Kamm, C., Fischer, H., Garavaglia, B., Kullmann, S., Sharma, M., Schrader, C., Grundmann, K., Klein, C., Borggraefe, I., Lobsien, E., et al. (2008). Susceptibility to DYT1 dystonia in European patients is modified by the D216H polymorphism. Neurology 70, 2261–2262.

Kim, C.E., Perez, A., Perkins, G., Ellisman, M.H., and Dauer, W.T. (2010). A molecular mechanism underlying the neural-specific defect in torsinA mutant mice. Proc Natl Acad Sci U S A 107, 9861–9866.

Kim, Y.S., and Leventhal, B.L. (2015). Genetic epidemiology and insights into interactive genetic and environmental effects in autism spectrum disorders. Biol Psychiatry 77, 66–74.

Kramer, P.L., Heiman, G.A., Gasser, T., Ozelius, L.J., de Leon, D., Brin, M.F., Burke, R.E., Hewett, J., Hunt, A.L., Moskowitz, C., et al. (1994). The DYT1 gene on 9q34 is responsible for most cases of early limb-onset idiopathic torsion dystonia in non-Jews. Am J Hum Genet 55, 468–475.

Lamming, D.W., Cummings, N.E., Rastelli, A.L., Gao, F., Cava, E., Bertozzi, B., Spelta, F., Pili, R., and Fontana, L. (2015). Restriction of dietary protein decreases mTORC1 in tumors and somatic tissues of a tumor-bearing mouse xenograft model. Oncotarget 6, 31233–31240.

Li, T.Y., Song, L., Sun, Y., Li, J., Yi, C., Lam, S.M., Xu, D., Zhou, L., Li, X., Yang, Y., et al. (2018). Tip60-mediated lipin 1 acetylation and ER translocation determine triacylglycerol synthesis rate. Nat Commun 9, 1916.

Liu, G.H., and Gerace, L. (2009). Sumoylation regulates nuclear localization of lipin-1alpha in neuronal cells. PLoS One 4, e7031.

MacVicar, T., Ohba, Y., Nolte, H., Mayer, F.C., Tatsuta, T., Sprenger, H.G., Lindner, B., Zhao, Y., Li, J., Bruns, C., et al. (2019). Lipid signalling drives proteolytic rewiring of mitochondria by YME1L. Nature 575, 361–365.

Martino, D., Gajos, A., Gallo, V., Cif, L., Coubes, P., Tinazzi, M., Schneider, S.A., Fiorio, M., Zorzi, G., Nardocci, N., et al. (2013). Extragenetic factors and clinical penetrance of DYT1 dystonia: an exploratory study. J Neurol 260, 1081–1086.

McNairn, A.J., Chuang, C.H., Bloom, J.C., Wallace, M.D., and Schimenti, J.C. (2019). Female-biased embryonic death from inflammation induced by genomic instability. Nature 567, 105–108.

Nestler, E.J., Pena, C.J., Kundakovic, M., Mitchell, A., and Akbarian, S. (2016). Epigenetic Basis of Mental Illness. Neuroscientist 22, 447–463.

O’Shaughnessy, P.J., Baker, P., Sohnius, U., Haavisto, A.M., Charlton, H.M., and Huhtaniemi, I. (1998). Fetal development of Leydig cell activity in the mouse is independent of pituitary gonadotroph function. Endocrinology 139, 1141–1146.

Ozelius, L., and Lubarr, N. (1993). DYT1 Early-Onset Isolated Dystonia. In GeneReviews((R)). M.P. Adam, H.H. Ardinger, R.A. Pagon, S.E. Wallace, L.J.H. Bean, K. Stephens, and A. Amemiya, eds. (Seattle (WA)).

Peterfy, M., Phan, J., Xu, P., and Reue, K. (2001). Lipodystrophy in the fld mouse results from mutation of a new gene encoding a nuclear protein, lipin. Nat Genet 27, 121–124.

Peterson, T.R., Sengupta, S.S., Harris, T.E., Carmack, A.E., Kang, S.A., Balderas, E., Guertin, D.A., Madden, K.L., Carpenter, A.E., Finck, B.N., et al. (2011). mTOR complex 1 regulates lipin 1 localization to control the SREBP pathway. Cell 146, 408–420.

Romani, P., Brian, I., Santinon, G., Pocaterra, A., Audano, M., Pedretti, S., Mathieu, S., Forcato, M., Bicciato, S., Manneville, J.B., et al. (2019). Extracellular matrix mechanical cues regulate lipid metabolism through Lipin-1 and SREBP. Nat Cell Biol 21, 338–347.

Sembongi, H., Miranda, M., Han, G.S., Fakas, S., Grimsey, N., Vendrell, J., Carman, G.M., and Siniossoglou, S. (2013). Distinct roles of the phosphatidate phosphatases lipin 1 and 2 during adipogenesis and lipid droplet biogenesis in 3T3-L1 cells. J Biol Chem 288, 34502–34513.

Shin, J.Y., Hernandez-Ono, A., Fedotova, T., Ostlund, C., Lee, M.J., Gibeley, S.B., Liang, C.C., Dauer, W.T., Ginsberg, H.N., and Worman, H.J. (2019). Nuclear envelope-localized torsinA-LAP1 complex regulates hepatic VLDL secretion and steatosis. The Journal of clinical investigation 130, 4885–4900.

Singh, S.M., McDonald, P., Murphy, B., and O’Reilly, R. (2004). Incidental neurodevelopmental episodes in the etiology of schizophrenia: an expanded model involving epigenetics and development. Clin Genet 65, 435–440.

Tanabe, L.M., Liang, C.C., and Dauer, W.T. (2016). Neuronal Nuclear Membrane Budding Occurs during a Developmental Window Modulated by Torsin Paralogs. Cell Rep 16, 3322–3333.

Tanabe, L.M., Martin, C., and Dauer, W.T. (2012). Genetic background modulates the phenotype of a mouse model of DYT1 dystonia. PLoS One 7, e32245.

Ulug, A.M., Vo, A., Argyelan, M., Tanabe, L., Schiffer, W.K., Dewey, S., Dauer, W.T., and Eidelberg, D. (2011). Cerebellothalamocortical pathway abnormalities in torsinA DYT1 knock-in mice. Proceedings of the National Academy of Sciences of the United States of America 108, 6638–6643.

vom Saal, F.S., and Bronson, F.H. (1980). Sexual characteristics of adult female mice are correlated with their blood testosterone levels during prenatal development. Science 208, 597–599.

Wang, X., Devaiah, S.P., Zhang, W., and Welti, R. (2006). Signaling functions of phosphatidic acid. Prog Lipid Res 45, 250–278.

Werner, R.J., Schultz, B.M., Huhn, J.M., Jelinek, J., Madzo, J., and Engel, N. (2017). Sex chromosomes drive gene expression and regulatory dimorphisms in mouse embryonic stem cells. Biol Sex Differ 8, 28.

Wijemanne, S., and Jankovic, J. (2015). Dopa-responsive dystonia--clinical and genetic heterogeneity. Nat Rev Neurol 11, 414–424.

Yang, C., Wang, X., Wang, J., Wang, X., Chen, W., Lu, N., Siniossoglou, S., Yao, Z., and Liu, K. (2020). Rewiring Neuronal Glycerolipid Metabolism Determines the Extent of Axon Regeneration. Neuron 105, 276–292 e275.

Yang, J.F., Wu, T., Li, J.Y., Li, Y.J., Zhang, Y.L., and Chan, P. (2009). DYT1 mutations in early onset primary torsion dystonia and Parkinson disease patients in Chinese populations. Neurosci Lett 450, 117–121.

Young, B.P., Shin, J.J., Orij, R., Chao, J.T., Li, S.C., Guan, X.L., Khong, A., Jan, E., Wenk, M.R., Prinz, W.A., et al. (2010). Phosphatidic acid is a pH biosensor that links membrane biogenesis to metabolism. Science 329, 1085–1088.

Zhao, C., Brown, R.S., Chase, A.R., Eisele, M.R., and Schlieker, C. (2013). Regulation of Torsin ATPases by LAP1 and LULL1. Proceedings of the National Academy of Sciences of the United States of America 110, E1545–1554.

Zhukovsky, M.A., Filograna, A., Luini, A., Corda, D., and Valente, C. (2019). Phosphatidic acid in membrane rearrangements. FEBS Lett.

