## Supplemental Videos for "Lipid metabolic stress in development defines which genetically-susceptible DYT-*TOR1A* mice develop disease"

### Slide 1
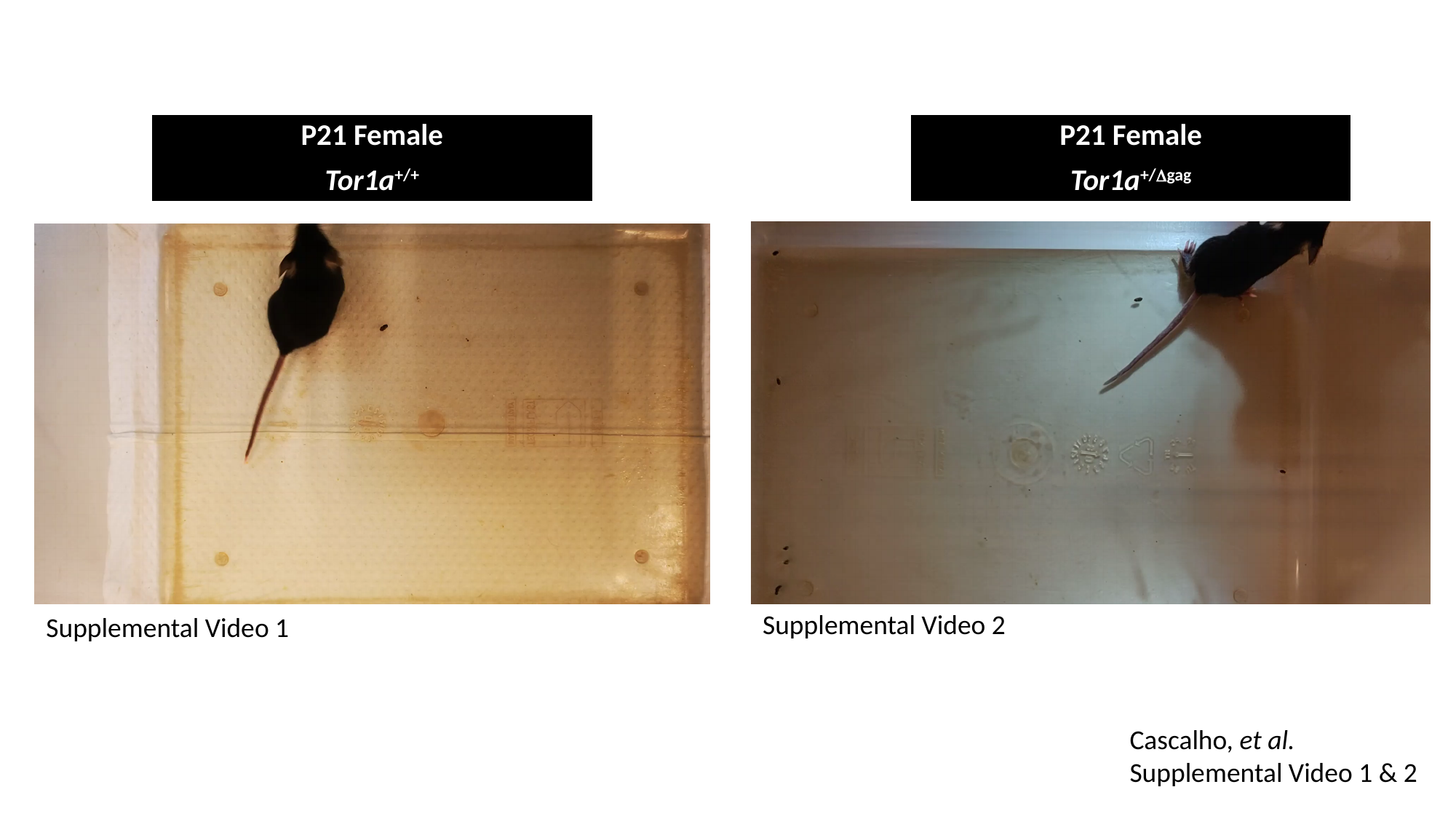

P21 Female
Tor1a^+/+^
P21 Female
Tor1a^+/Δgag^
Supplemental Video 2
Supplemental Video 1
Cascalho, et al.
Supplemental Video 1 & 2
